# Droplet-assisted folding of long regulatory RNAs

**DOI:** 10.1101/2025.09.15.676367

**Authors:** Simon Doll, Lukas Pekarek, Fathima Ferosh, Jovana Vasiljević, Tyler Harmon, Marcus Jahnel

## Abstract

Long regulatory RNA regions orchestrate complex cellular processes, including gene expression and epigenetic modifications. How these RNAs dynamically fold and refold in response to cellular signals remains poorly understood. Given that RNAs interact with ubiquitous RNA-binding proteins (RBPs) prone to form biomolecular condensates, we explore how protein droplets interacting along an RNA impact its folding process. Attached droplets prevent premature folding by competing with RNA:RNA interactions. When droplets dissolve due to cellular signals, capillary effects cause the RNA to collapse while refolding. We test this process of condensate-guided RNA folding by adapting established RNA secondary structure predictors to mimic various folding pathways and supplement this with coarse-grained simulations. We find that interactions with transient droplets robustly leads to the formation of long-range RNA contacts, which are otherwise hard to achieve. Our results compare favorably with available experimental data. We propose that this strategy, which we call droplet-assisted RNA folding, represents a previously unexplored mechanism for shaping RNA structures. Given the widespread propensity of RBPs to form condensates, this process could play a fundamental role in the structural organization, conditional reshaping, and functional regulation of long regulatory RNAs.

**Significance:** Complex non-coding RNA regions (e.g. lncRNAs, 3’UTRs, etc.) perform several important functions in higher organisms. However, the longer an RNA, the more likely it is to misfold. How can such regulatory regions of more than 1000 nts be reliably folded, considering that premature co-transcriptional folding favors local interactions? We present a mechanism by which biomolecular condensates of RNA-binding proteins (RBPs) assist and control the folding of complex RNAs. Here, droplets of RBPs nucleate around RNA binding motifs, prevent premature misfolding through their RNA chaperone function, and allow robust folding of initially distant regions through coordinated droplet dissolution and detachment. This mechanism provides an alternative perspective on how long RNAs acquire their structures in a context that is functionally dynamic.

## Introduction

The functional versatility of RNA underscores the crucial role of RNA structures [1–3]. Proper folding enables distant segments of an RNA molecule to come into close proximity, facilitating essential biological functions [4–6]. However, some RNA species-such as mRNAs, rRNAs, and lncRNAs-can be over 1000 nucleotides (nt) long [7]. This raises a fundamental challenge: how do living organisms ensure the robust and accurate folding of long regulatory RNAs [8]?

Efforts to understand RNA folding have led to the development of numerous RNA structure prediction algorithms [9–13]. Among them, thermodynamic-based minimum free energy (MFE) models have become the gold standard. These models predict the most stable RNA structure by identifying the global minimum in free energy, without identifying a mechanistic pathway of the folding process. However, the most thermodynamically stable structure does not necessarily correspond to the functional conformation — RNA structures are inherently dynamic [1–3].

Inspired by the vectorial 5’ to 3’ nature of transcription, co-transcriptional folding algorithms aim to capture the stepwise formation of RNA structures as they emerge from RNA polymerase [11, 14, 15]. These approaches not only improve structure prediction accuracy, but also provide mechanistic insights into RNA folding. However, certain experimental observations challenge the explanatory power of co-transcriptional folding alone. For instance: (**1**) the ends of long RNAs are often found in close proximity despite their linear separation [4, 6], (**2**) downstream sequences can induce RNA refolding *in vivo* but not *in vitro* [16], and (**3**) the same RNA can adopt different conformations depending on cellular conditions [17]. These findings suggest that transcription-driven folding alone does not fully dictate the final RNA structure.

Within cells, RNAs are constantly surrounded by a multitude of RNA-binding proteins (RBPs) [18–20]. RBPs, which are among the most intrinsically disordered protein classes [21, 22], frequently form dynamic, liquid-like molecular assemblies known as biomolecular condensates [19, 20, 22, 23]. These condensates are implicated in key biological processes such as transcription regulation, RNA splicing, and stress responses in eukaryotic cells [24–28].

While RBP interactions with RNA are often sequence-dependent, protein binding can also induce RNA structural rearrangements or even cause structural dissolution [29–32]. The engulfment of RNAs within biomolecular condensates could provide a mechanistic basis for the long-range interactions observed in various experimental studies [5, 33–37]. Furthermore, biomolecular condensates, similar to other systems studied in soft matter physics [38], have been shown to exert capillary forces on biopolymers and membranes [39, 40], providing the potential to influence RNA folding dynamics.

Importantly, the formation of these condensates is tightly regulated by factors including temperature, pH, protein and RNA composition, and concentration, as well as post-translational modifications [20, 41–46]. Prominent RNA-binding proteins like G3BP1, FUS or YBX1 can be post-translationally modified distinctly in either the RNA-binding domain or in the IDR [47–49].

For example, ubiquitination of G3BP1 prevents the phase separation of the protein and stress granule formation, while acetylation K376 of the RRM RNA binding domain prevents both RNA binding and phase separation [47, 50, 51]. Another example is YBX1, which can be phosphorylated by the kinase Akt1 at Ser102 to modulate RNA binding or in the IDR by FGFR1 kinase [52], potentially affecting its phase behavior. Interestingly, post-translational modifications (PTMs), such as methylation or phosphorylation of the SARS-CoV-2 Nucleocapsid, have also been shown to regulate both the RNA-binding propensity and the material properties of the formed condensates [41, 53, 54].

Motivated by these observations, we propose an alternative scenario — **Droplet-assisted RNA folding** — in which droplets of biomolecular condensates shape RNA structures. The key elements of this scenario are based on the following findings/principles: (i) The ability of droplets to shape long polymers through capillary forces [55], (ii) The propensity of RBPs to form biomolecular condensates (droplets), and (iii) The regulation of the formation of biomolecular condesates in cells through PTMs. Here, we mimic droplet-assissted and other folding scenarios through the differential incorporation of constraints into established MFE-based secondary structure prediction algorithms. Additionally, we supplement this with coarse-grained simulations of RNA folding in the presence of a transient droplet. Our approach compares favorably with experimental structures from the PDB. Finally, we apply this concept to two widely studied long RNAs, the lncRNA HOTAIR and the SARS-CoV-2 genomic RNA. The resulting, deeply nested structures are characterized by long-range interactions and are unique in comparison with published results, indicating that a droplet-assisted folding process provides new and biologically motivated insight into the folding pathway of long RNAs.

## Results

### Guiding RNA structure formation by different RNA folding scenarios

To incorporate the influence of condensates and mimic droplet-guided RNA folding, we decided to employ well-established RNA folding algorithms ViennaRNA [9] and RNAstructure [10] to guide the folding process using strategic constraints (**See Methods**). In detail, we iteratively update base pairing restrictions according to the different scenarios (**Fig. 1**). Besides the common, restriction-free **Full** (minimum free energy folding), we implemented a co-transcriptional approach, here termed **Sequential** by folding the RNA step-wise from the 5’ to the 3’ end. For the **Single Droplet** approach, the RNA is allowed to fold symmetrically, starting from both ends. In the **Multiple Droplets** scenario, a number of droplets are distributed over the length of the RNA. Initially covering the whole sequence, these droplets shrink simultaneously in a given number of dissolution steps, allowing the exposed nucleotides to form base pairs. In this study we always implemented 4 droplets, equally spaced along the given RNA sequence.

**Figure 1.**
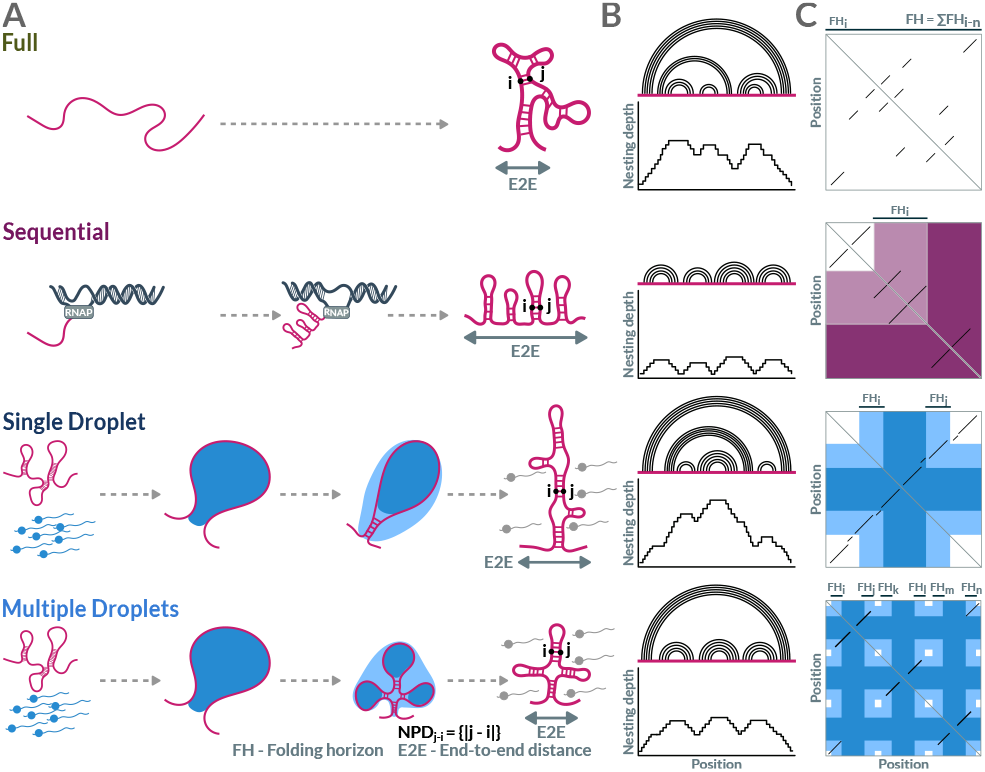
Various RNA folding pathways can lead to distinct structural features. **A**) Schematic illustration of different RNA folding scenarios. The analysed parameters (End-to-end distance (E2E) and Nucleotide pair distance (NPD)) are outlined . **B**) Arc diagrams and plots of the nesting depths to illustrate the conceptual idea of expected structures resulting from the different folding scenarios. **C**) Conceptual heat maps and illustration of the folding horizon (FH) for each scenario. Darker purple or blue areas of interactions only become accessible at later steps of the folding process.

We hypothesize, that by applying different folding scenarios it is possible to robustly obtain distinct RNA structures, defining a framework for their dynamic re-shaping. Thus, structures obtained during transcription might change after interacting with a condensate.

### Folding pathways robustly shape RNAs in a sequence-independent manner

To compare and characterise the resulting structures we calculated multiple parameters and distributions for all predictions: the minimum free energy (MFE) as a characteristic of a structure’s stability; the end-to-end distance (E2E), defined as the number of nucleotides in the exterior loop [6]; the nesting depth of each nucleotide, which is the cumulative sum of unclosed base-pairs from the 5’ send; and the cumulative distribution of the distances between each base-paired nucleotides (NPD, normalized to the RNA sequence length) (**Fig. 2B-E**).

**Figure 2.**
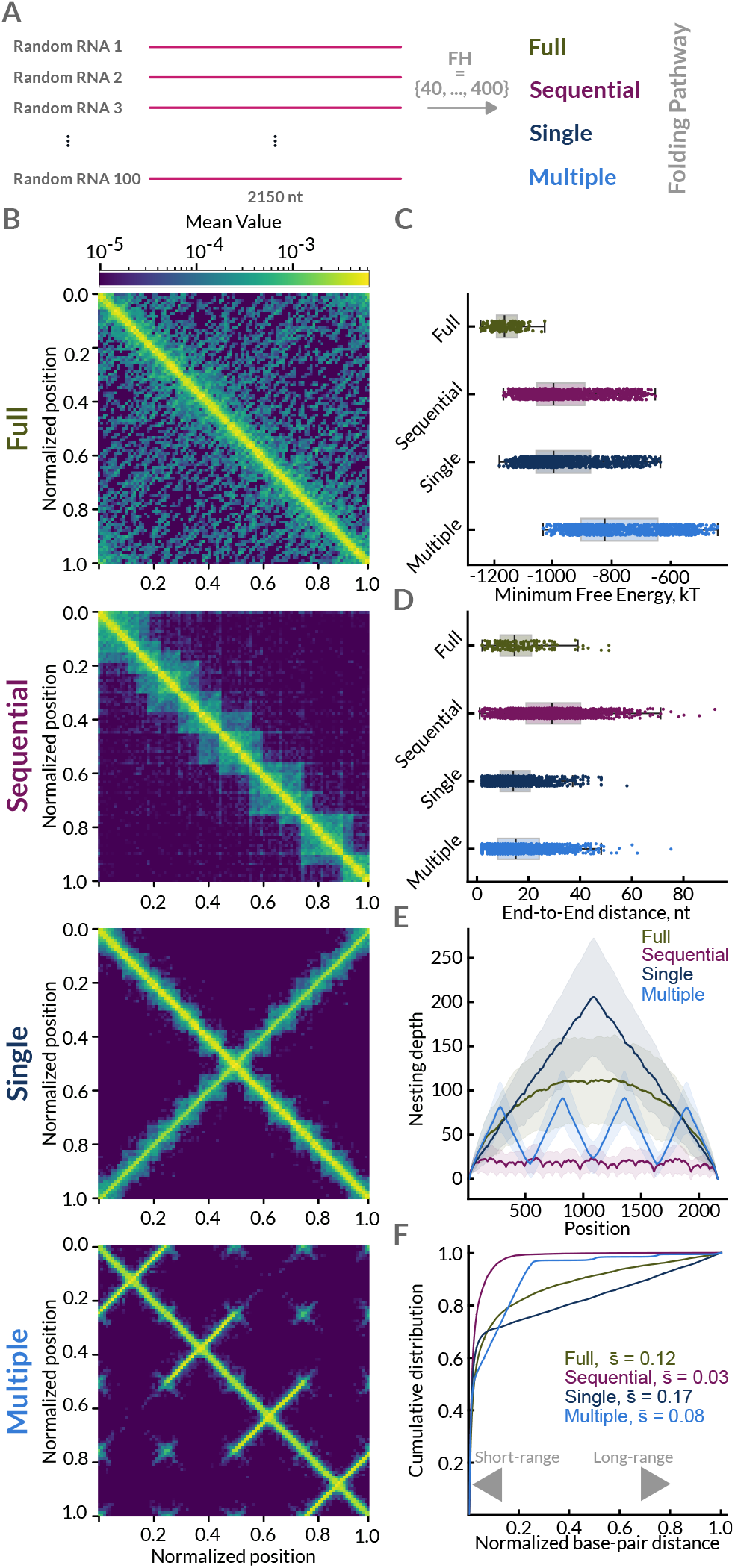
RNA folding scenarios robustly shape randomized RNAs. **A**) Results for structure predictions for 100 random RNAs for different folding scenarios. **B**) Coarse heat maps of cumulative base-pairing matrices across different folding horizons. **C-F**) Distributions of different structure parameters: **C**) Minimum Free Energy **D**) End-to-End distance **E**) Nesting depth (solid line representing the average; the shaded area represents the 95% confidence interval obtained via bootstrapping) **F**) Cumulative distribution of normalized base-pair distances; normalized to the length of the RNA. 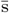 signifies the mean normalized base pair distance.

To test the robustness of the folding scenarios, we generated 100 random long RNAs of equal length (2150 nt, inspired by lncRNA HOTAIR) and performed all four different folding pathway predictions over various folding horizons. We define the folding horizon (FH) as the number of nucleotides becoming additionally available in each iterative step. The different folding pathways yielded RNA structures of distinct patterns, as shown in the average contact heatmaps (**Fig. 2B**). The **Full** prediction comprised many randomly dispersed interactions, with the 5’ and 3’ ends predicted to base-pair with each other. **Sequential** predictions favor local, short-range interactions with hairpin size depending on the chosen folding horizon. **Single Droplet** folding resulted in a deeply nested structure (indicated by the perpendicular diagonal in the corresponding heatmap). Small substructures were predicted to occur for longer folding horizons, but the overall nature of deeply nested structures remained in all predictions. Finally, the **Multiple Droplets** pathway yielded structures with characteristics between the short-range **Sequential** predictions and the deeply nested structure of the **Single Droplet** scenario. Importantly, this in-between pathway robustly predicts the formation of long-range interactions between distant regions, including the 5’ and 3’ ends. Consequently, the **Multiple Droplet** pathway leads to the formation of nested structures located at seed positions.

The **Full** predictions showed the lowest and most narrowly distributed MFE values, with the other pathways yielding more diverse MFE distributions (**Fig. 2C**). As expected, the **Sequential** folding predictions possessed the most extended E2E distance (**Fig. 2D**). The E2E distribution of **Full, Single Droplet** and **Multiple Droplets** showed no significant difference between each other. The nesting depth of the **Sequential** folding predictions show the lowest values for all nucleotides, with frequent opening and closing of small hairpin structures (**Fig. 2E**). The **Full** predictions nesting depth resemble a semicircle distribution with the largest uncertainty. This signature of freely independent probabilities is a result of the randomly generated RNAs, which can also be seen in the corresponding contact map. The nesting depths of the **Single Droplet** and **Multiple Droplets** predictions agree with the assumed behaviour: The **Single Droplet** predictions have the highest nesting depth in the middle of the sequence, with a continuous increase before and a continuous decrease afterwards. The **Multiple Droplets** predictions have four characteristic peaks in the nesting depth at the position of the seeds and while there are local minima in-between them, the nesting depth is not decreasing to zero, indicating a long-range interactions between distant parts of the RNA (**Fig. 2E**).

Finally, we calculated the cumulative distribution of all the distances between base-paired nucleotides (NPD) to obtain an overall descriptor of the predicted structures. This characterizes global base-pairing differences between structures of distinct folding pathways, such as preferences for short- or long-range interactions (**Fig. 2F**). While **Sequential** structures in this measure again show predominantly short-range interactions, the structures predicted following the **Single Droplet** pathway show the highest tendency for long-range interactions. The **Full** and **Multiple Droplets** structures show an NPD between these extremes, with the **Full** structures exhibiting more long-range interactions. Taken together, these calculations indicate that the folding pathway robustly determines the global organisation of a long RNA.

### Multiple droplets pathway captures global characteristics of experimentally determined RNAs

Next, we wondered whether we could guide structure formation of native RNAs that evolved to have specific shapes and functions. To this end, we extracted representative 3D structures of long RNAs from the RCSB PDB database (**see methods**) [56]. This allowed us to evaluate the predictions against a “ground truth” of experimentally determined RNA structures (**Fig. 3**).

**Figure 3.**
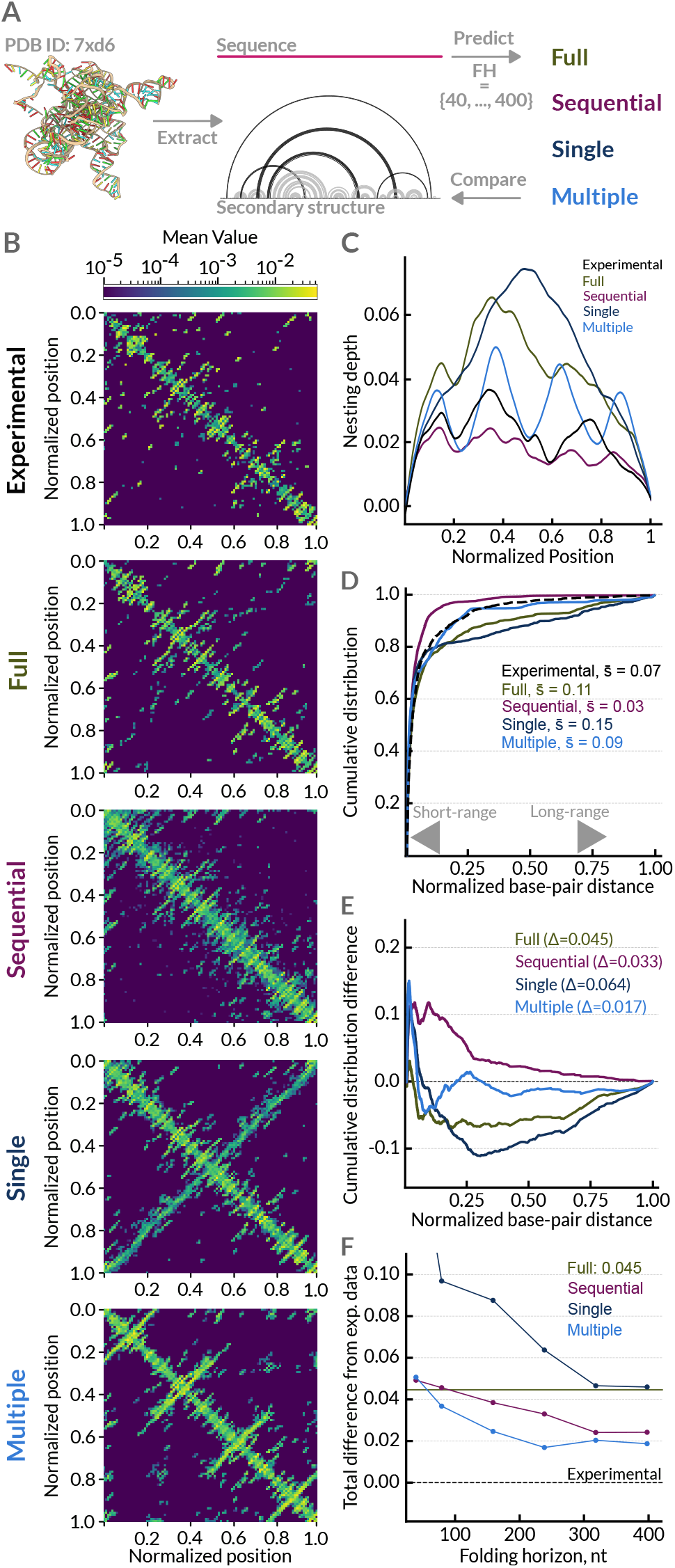
Droplet-assisted folding pathways capture characteristics of experimentally resolved RNA structures. **A**) Comparison of selected experimentally resolved RNA structures (obtained from PDB database) with predicted structures for different folding scenarios. **B**) Coarse heatmaps of cumulative base-pairing matrices across different folding horizons. **C-F**) Distribution of different structure parameters: **C**) Nesting depth **D**) Cumulative distribution of normalized base-pair distances. 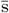 signifies the mean normalized base pair distance.**E**) Difference between distributions of normalized base-pair distance for experimental data and folding scenarios. The total difference (calculated as the area between the curves) is shown in brackets. Data shown in **D** and **E** is for FH = 240 nt. **F**) Dependence of the total difference of the cumulative distribution from the experimental distribution on the folding horizon size. Normalizations are to the length of the RNA.

To compare the folding process on RNAs of different lengths, we normalized and averaged the contact maps and other parameters (MFE, E2E, etc.). The different folding pathways yielded distinct contact map patterns similar to those observed for the random RNAs, albeit the patterns were not as emphasized. For example the characteristic cross-shape of the **Single Droplet** pathway is again apparent (**Fig. 3B**). The different folding scenarios did not yield any significant difference in the normalized MFE values, and the E2E values of the experimentally determined structures were larger than all the other folding pathways, except the **Sequential** one (**Fig. S3** and **Table S1**).

The nesting depth of the **Sequential** predictions is again the lowest one for all nucleotides, although single hairpins are not as visible as before. Also, the **Single** and **Multiple Droplet** nesting depths follow a similar trend as observed before, although with less pronounced features. Interestingly, the **Full** prediction nesting depth still shows a resemblance to a semicircle distribution, indicating a certain randomness in predicting such large structures. In comparison to these four, the nesting depth of the experimental structures show the second lowest nesting depth for each nucleotide, only higher than the **Sequential** predictions, with some substructures being distinguishable (**Fig. 3C**).

Finally, we examined the distribution of NPD (**Fig. 3D**). Here, the distribution curve of the **Multiple Droplets** predictions closely follows the distribution of the experimental data. To verify this observation, we calculated the difference between the cumulative distributions of experimental data and each folding scenario (**Fig. 3E**). In addition, we explored the effect of the folding horizon on the difference between the experimental and predicted NPD distributions. Again, the **Multiple Droplets** pathway performs best in all tested cases with an optimum around a 400 nt folding horizon (**Fig. 3F**).

Taken together, the comparison of experimental data with different folding pathways indicates that a folding process guided by dissolving condensate droplets can explain global characteristics of known long RNA structures.

### RBP YBX1 and lncRNA HOTAIR as a model system

So far, to follow the general idea of RNA folding as a result of a dissolving droplet losing contact, we used model parameters (folding horizon, number and position of seeds) without reference to any biological system. Although some RBPs bind unspecifically, many have known binding motifs or preferences for certain sequences [19]. These local differences give rise to a global variable binding probability across a given RNA sequence which can result in uneven droplet dissolution from the RNA. By using available data in the literature to predict the binding preferences of a given RBP along a given RNA sequence, we were able to guide the RNA folding process in a more biologically relevant manner. For example, YBX1 is an abundant RBP that features a conserved cold-shock domain flanked by extended IDRs to contact RNA at specific RNA sequences [56, 57]. One of the many RNAs YBX1 has been shown to interact with is the long non-coding RNA (lncRNA) HOTAIR [58]. As lncRNAs do rarely undergo translation, and thus no periodic unfolding of the RNA structures, we use HOTAIR-YBX1 as our model system to study droplet-assisted folding in a biologically relevant scenario. We first confirmed the propensity of YBX1 to form biomolecular condensates and co-localize with HOTAIR by confocal microscopy experiments in vitro (**Fig. 4A** and **Fig. S4A**).

**Figure 4.**
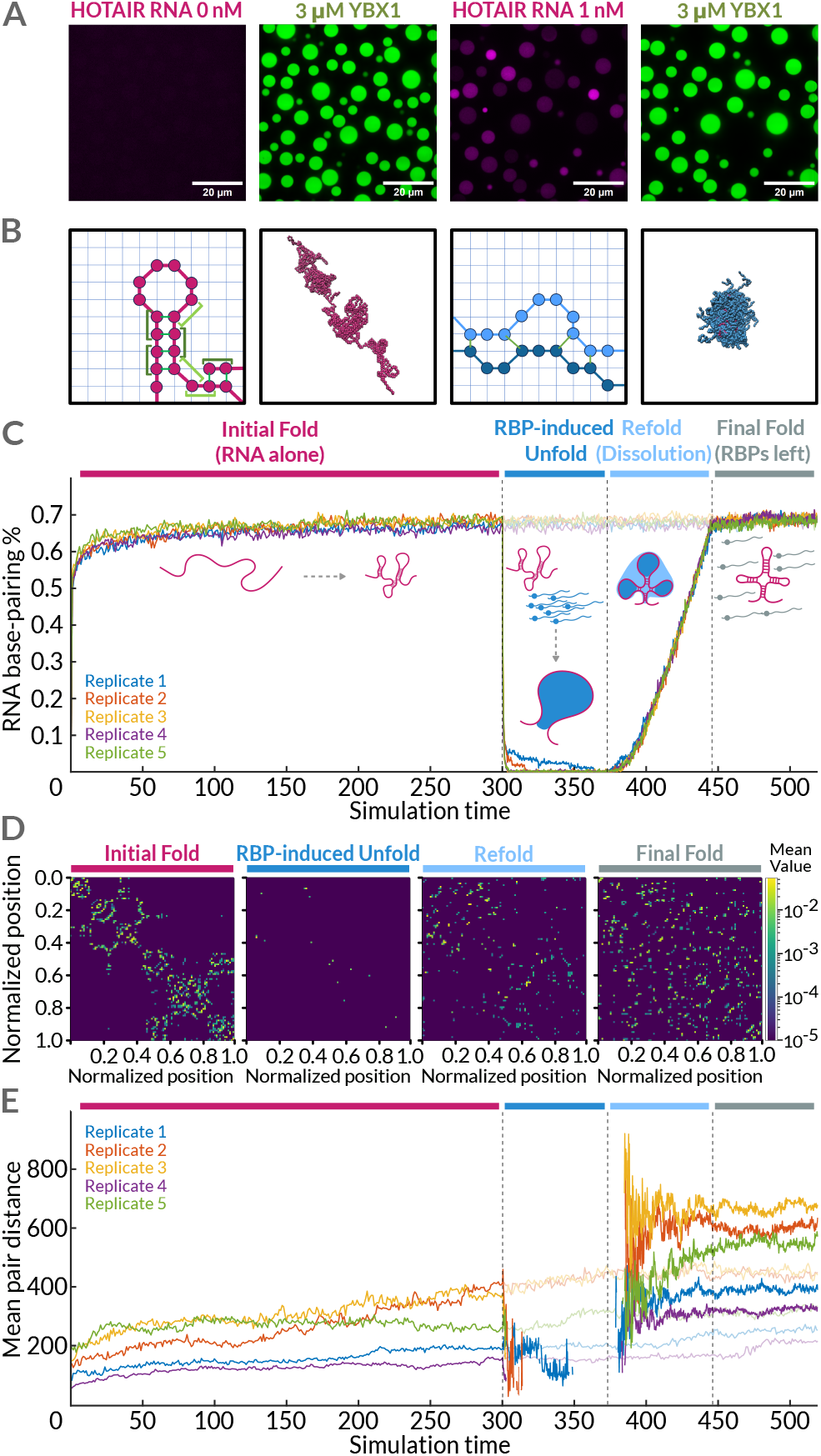
Droplet-assisted folding increases long-range interactions in simulations. **A**) Confocal fluorescence microscopy image of co-condensation of RBP YBX1 without (left side) and with lncRNA HOTAIR (right side). HOTAIR strongly partitions into YBX1 droplets. YBX1 was fused with EGFP, HOTAIR RNA was internally labeled with modified UTPs. **B**) Schematics and representative snapshots of lattice-based simulations of RNA (pink) and protein (blue). Each bead represents a nucleotide or two amino acids with each lattice site corresponding to *∼*8 Å. Green lines show stoichiometric interactions for both RNA and protein and green brackets show nearest neighbor contributions for RNA base stacking (dark green) and stem ends (light green). Snapshots are zoomed in from a 3D periodic lattice of length 333. **C**) Fraction of RNA base pairs over time. Each color corresponds to an independent simulation. The corresponding muted lines shows the folding process for simulations where no droplet had been added. **D**) Representative contact maps at different points along the simulation trajectory - end of initial fold, end of unfold, end of refold, and the final conformation. The transient droplet causes a global reorganization in contacts which promotes long-range interactions seen as contacts far from the diagonal. **E**) Mean nucleotide distance between contacts along the simulation trajectories. Muted lines show the simulations without a droplet. Note, during the unfold phase where there are near-zero RNA:RNA interactions this metric is ill-defined resulting in a gap in the data. A large increase in the mean contact distance occurs during refolding from droplet dissolution because the RNA structures became single stranded and collapsed. The rank-order persists in the simulations because the RNA dynamics are slow compared to the short lifetime of the droplet.

### Coarse-grained simulations demonstrate the dynamic reshaping of long RNAs through condensate droplets

To demonstrate that droplets of RNA-binding proteins could cause large-scale reorganizations in the resulting folded state, we turned to coarse-grained simulations. We used a lattice polymer model with one-to-one stoichiometric interactions that has been successful in modeling phase separated proteins [57–59] which was coarse-grained to 1 nucleotide or 2 amino acids per bead (*∼* 8 Å); see cartoon and snapshots in Figure (**Fig. 4A**). We extended the model by adding RNA-like molecules with stoichiometric interactions for base pairing, nearest-neighbor interactions for base stacking and antiparallel structure, and geometry constraints for loop penalties. We chose protein:protein interactions that resemble the architecture of YBX1 and phase separates at *∼*1 *µ*M, we chose RNA:RNA interactions consistent with lncRNA HOTAIR that match the energy tables of the Nearest Neighbor Database [60, 61], and chose strong protein:RNA interactions which are consistent with 1 nM binding affinity (measured by fluorescence anisotropy, data not shown); see SI for further details on the simulations.

The simulations (5 replicas) were performed in four stages as illustrated in the schematic **Fig. 4B. Initial Fold (RNA alone):** First the RNA was allowed to fold with no proteins present (3 × 10^11^ steps, 1.4 × 10^8^ steps per bead). **RBP-induced Unfold:** Then 50 RNA-binding proteins (5*µ*M) were added in close proximity to the RNA and allowed to interact (3 × 10^11^ steps, 3.4 × 10^7^ steps per bead). **Refold (Droplet dissolution):** Next the proteins were one-by-one converted into a weakly interacting protein that disassociates from the droplet to mimic phosphorylation induced disollution and RNA detachment (6 × 10^9^ 50 steps, 6.8 × 10^5^ 50 steps per bead). **Final Fold (RBPs left):** Finally, the RNA was allowed to equilibrate with the 50 weakly interacting proteins present (3 × 10^11^ steps, 3.4 ×10^7^ steps per bead). For later comparisons, the simulations were duplicated after the initial folding step. One set of simulations experienced the droplet as described above, the other continued folding with no proteins ever added (6×10^11^ steps, 2.8 × 10^8^ steps per bead).

**Figure 4C-E** quantify how the simulations behaved through the cycle of adding and dissolving a droplet with the reference no droplet simulations shown as shadows. **Figure 4C** shows the fraction of base-pairing in the simulations. The initial fold yields a rapid convergence to high base-pairing with little further increase. After undergoing a droplet cycle in which the stems were dissolved, the fraction rapidly converges to the no-droplet base-pairing fraction, almost entirely limited by the simulation detachment rate. **Figure 4D** shows the contact maps at different points along the trajectory for a representative simulation. It highlights that the RNA structure changes from having primarily local interactions to one that has primarily global interactions. We quantify this in **Figure 4E** which shows the average nucleotide distance between existing base pairs. The initial fold locks local structure in with large energy barriers to reorganization showing interactions closer to the diagonal. When refolded during droplet dissolution, the RNA is in a more collapsed state, where distal parts of the RNA can lock in a much more global structure where stems form much farther off the diagonal. Together, our simulations support the notion that droplets can dramatically alter the folding pathways realized by long RNA.

### Information about RNA-binding motifs can guide the predictions

The interactions between RNA-binding proteins and RNAs have been extensively studied [62, 63]. While some RBPs bind unspecifically, many have known binding motifs or sequence preferences [19]. These local differences result in a variable global binding probability across a given RNA sequence. As many RBPs are prone to form condensates, we reason that this variable binding profile can lead to an uneven RBP droplet detachment processes from the RNA. Using publicly available data, we next predict the binding preferences of an RBP along an RNA sequence in order to guide the RNA folding process in a more biologically relevant manner.

To this end, we extracted experimentally determined binding motives for YBX1 from the ATtRACT database [64, 65] and calculated the binding density of YBX1 along the full HOTAIR sequence (**Fig. 5A**). Next, we used this irregular binding density distribution as the base for our droplet dissolution predictions. We call this the **RBPFold** pathway. We tested different numbers of dissolution steps, ranging from 5 to 15 (**Fig. S4C**), where more steps correspond to a slower dissolution process (implemented as a smaller folding horizon). Finally, we compare our predictions with an experimentally determined HOTAIR structure based on chemical probing [66]. We extracted this secondary structure and analyzed it similarly to our predictions (**Fig. B-D** and **Fig. S4**).

**Figure 5.**
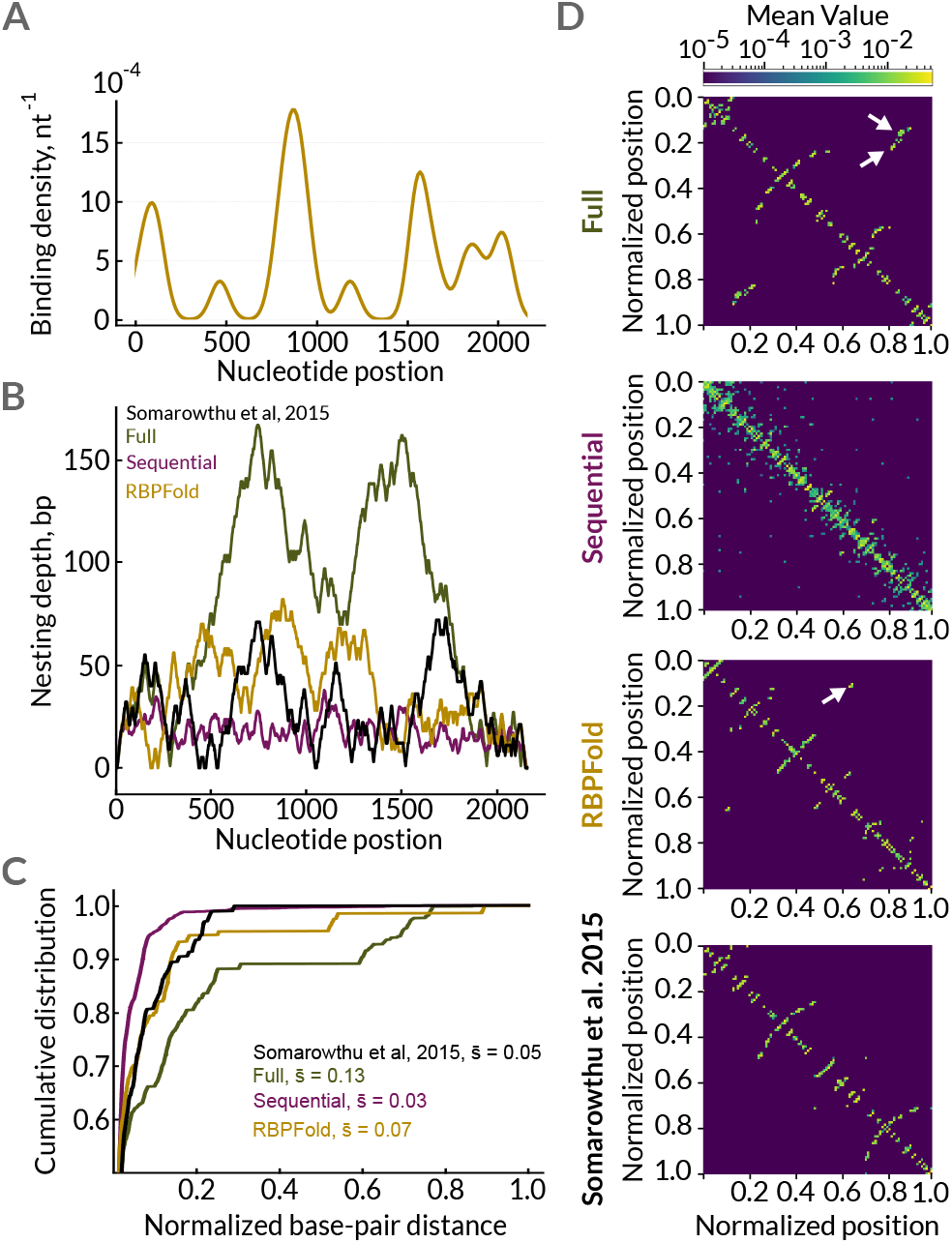
YBX1 and lncRNA HOTAIR illustrate droplet-guided folding process in a biological context. **A**) Calculated binding profile of YBX1 along HOTAIR shows prominent peaks and valleys. **B**) Nesting depths of multiple predicted and the proposed structure by Somarowthu *et al*. over the nucleotide position. **C**) Cumulative distribution of base-pair distances (normalized by RNA length) indicates increased fraction of long-range interactions with RBPFold over the experimental structure. **D**) Coarse heatmaps of cumulative base-pairing matrices for different folding predictions. The structure predicted for the RBPFold pathway are calculated for 15 binding levels. Arrows highlight long-range interactions.

The *in vitro* sample preparation by Somarowthu *et al*. 2015 reflects the co-transcriptional folding of HOTAIR [66], without any further refolding process (e.g. heating-cooling cycles or interaction with proteins). This is reflected in our analysis: the experimental E2E distance is even longer than the **Sequential** predictions (**Fig. S4E**) and with commonly higher nesting depths than the **Sequential** predictions, similar to the **RBPFold** and lower than the **Full** (**Fig. 5B**). Moreover, the NPD distribution of the Somarowthu data has the second strongest tendency for short distance base-pairing, only lower than the Sequential ones (**Fig. 5C**).

The **RBPFold** resulted in less stable (higher MFE) but more packed (lower E2E distance) structures (**Fig. S4D-** F). The nesting depth and the NPD distribution show a higher tendency to long-range interactions, though not as dominant as for the **Full** prediction (**Fig. 4D**, indicated by arrows). Thus, by combining binding predictions and available protein binding information, we can fine-tune the droplet-guided predictions of RNA structure formation in a biologically rationalized manner.

### Binding-guided structures of the SARS-CoV-2 RNA genome and Nucleocapsid protein mimic experimental cross-linking data

Within eukaryotic cells, RNAs undergo various processes, ranging from transcription, splicing, and translation to stress granule formation. This inevitably means that RNAs interact with many different RBPs [18–20]. While our approach, in principle, allows for the simulation of heterogenous droplets formed by multiple RBPs, we decided to seek a more isolated RNA-RBP system in which the interaction would be, at least at some point in the RNA lifetime, more defined and exclusive. Viral RNAs represent an example of such a system. Viral genomes often consist of one or more long RNA molecules that, at least within the virion particle, have one or a maximum of a few interaction partners. Analogous to host RBPs, viral RNA-binding proteins often contain IDRs and induce condensate formation [67–70]. Here, the SARS-CoV-2 RNA genome and the cognate Nucleocapsid protein (N) suit as a test pair for our method. The genomic RNA of SARS-CoV-2 is almost 30,000 nt long, yet within the virion particle, it interacts almost exclusively with the N protein thus forming the nucleocapsid of the virus. Moreover, due to the recent pandemic, SARS-CoV-2 genomic RNA is among the most thoroughly studied long RNAs to this date. Although no 3D structural data is available for the full genomic RNA, there are multiple chemical probing studies [71–73]. Several studies also performed RNA cross-linking, allowing them to detect even long-range interactions within the RNA directly [74–76].

We calculated the binding density of the N protein along the SARS-CoV-2 RNA genome based on the published data by Fan *et al*. 2022 [77](See **methods, Fig. 6B**). We then predicted RNA structures according to the different folding pathways together with the binding-guided RBPFold. Additionally, we combined the binding data with the chemical probing data from [72, 73] as an additional input, yielding a **RBPFold + Probing** folding pathway, which reflects both the structure shaping by a known RBP interactor and the experimentally determined base-pairing probabilities (**Fig. S5**). We also compared our data with the structure predicted by Lan *et al*. 2022 and Huston *et al*. 2021. Of note, the chemical probing-inspired structure predictions are often generated with the predefined restriction on maximum pairing distance [72, 73], therefore, these predicted structures likely underestimate possible long-range interactions.

**Figure 6.**
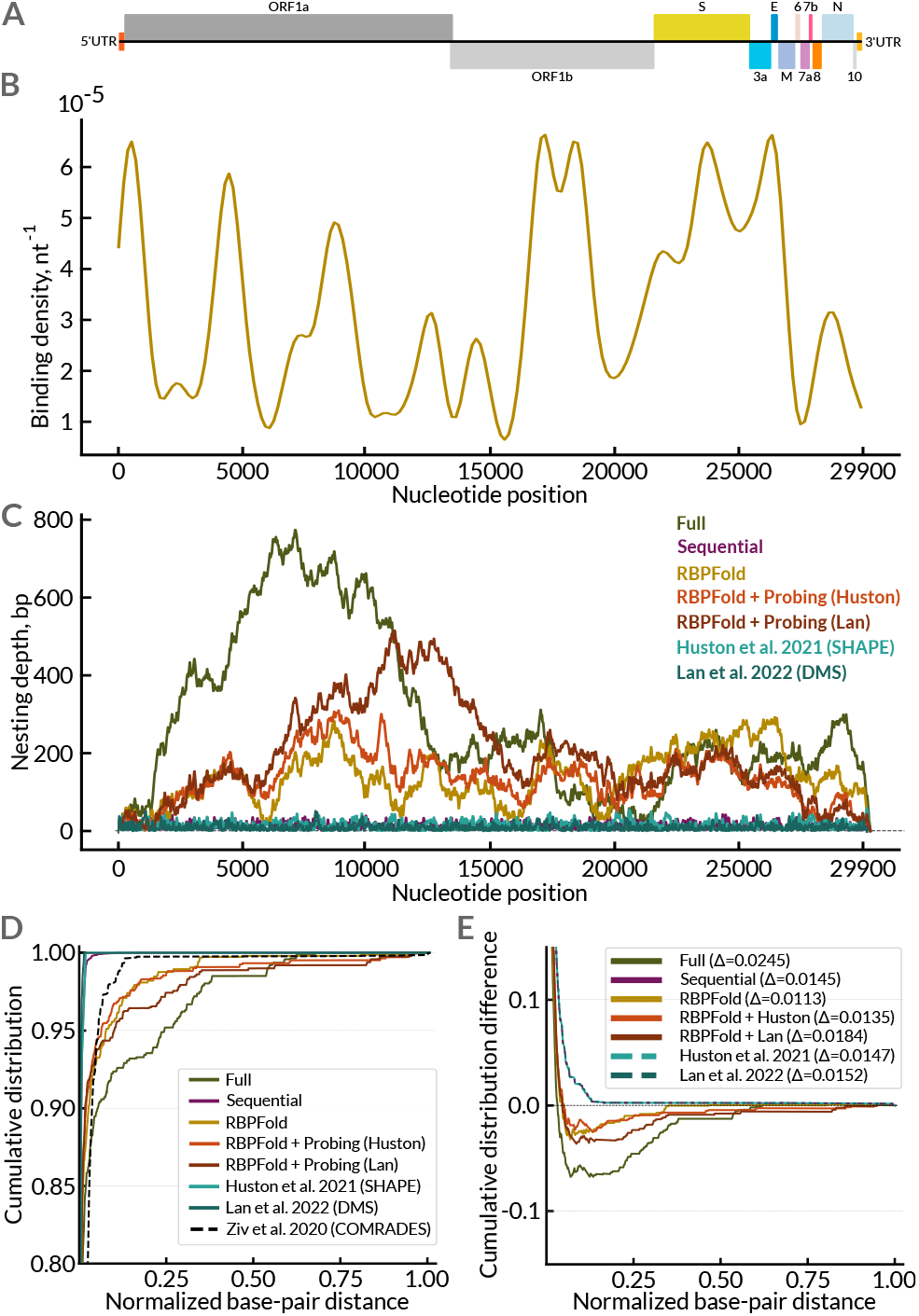
SARS-CoV-2 genomic RNA & Nucleocapsid protein. **A** Depiction of genomic regions of SARS-CoV-2 and **B**) binding profile of the nucleocapsid protein along the RNA. **C**) Nesting depth of predicted and an experimentally determined structure over the nucleotide position. **D**) Cumulative distribution of normalized base-pair distances for different folding scenarios and experimentally informed structures. **E**) Difference between the normalized base-pair distance distributions between the shown structures and COMRADES data [34]. The total difference (calculated as the area between the curves) is shown in brackets.

To analyse the resulting structures, we focused on the NPD distributions and compared them with the *in vivo* cross-linking data provided by Ziv et al. 2020 [34]. As mentioned above, the cross-linking and RNA chimeras sequencing experiments provide an insight into RNA interactions, including the long-range contacts. Interestingly, the NPD distribution of the cross-linking data is located between the short-range-favoring Sequential and chemical probing-inspired data [72, 73] and the long-range-allowing droplet-inspired predictions (**Fig. 6D** and **Fig. S5B**).

We then took the cross-linking data as a proxy of the ground truth and calculated the difference distributions (**Fig. 6E** and **Fig. S5C**). Again, RBPFold and RBPFold+Probing predictions are very close to the experimental base-pairing distribution. By taking into account the RNA sequence, the irregular binding profile of an RBP, and the chemical probing data, our droplet-guided folding pathway provides a biologically rationalized framework, in which deeply nested structures and long-range interactions arise naturally. The predicted secondary structures constitute a new class of RNA shapes that closely match experimental data.

## Discussion

Originally, it was assumed that the structure of a protein is determined by sequence alone. In analogy to protein folding, it had been traditionally assumed that an RNA’s sequence is the main factor determining its structure. However, it was shown that proteins, especially complex ones, require an orchestrated co-translational folding [78, 79] and additional chaperone factors [80] to fold correctly. Recently, researchers have been considering co-transcriptional folding as a biologically inspired folding pathway for RNAs [81]. Our motivation is to consider the role of RNA chaperones in folding and refolding of long RNAs.

RNA has unique physical properties compared to proteins. The promiscuous nature of base-pairing in a limited alphabet leads to a rugged energy landscape for folding with a high degree of degeneracy [1]. This leads to the folding process of a long RNA to be predominantly determined by which nucleotides meet first, establishing the first stable cornerstones around which the structure emerges. Additionally, the chaperones for RNA folding are RBPs, many of which have been shown to phase separate – which invokes the physics of different chemical phases as well as capillary and surface effects.

To explore this, we introduce a new model of RNA folding: **Droplet-Assisted Folding** (**Fig. 7A**). In this model, droplets of RNA-binding proteins disrupt RNA:RNA interactions, allowing for a collapsed, single-stranded, and unfolded RNA ensemble. As the droplets dissolve and detach due to posttranslational modifications, the balance of protein and RNA interactions shifts towards RNA:RNA interactions, which allows new RNA stems (the cornerstones) and subsequent folding from this unique state.

**Figure 7.**
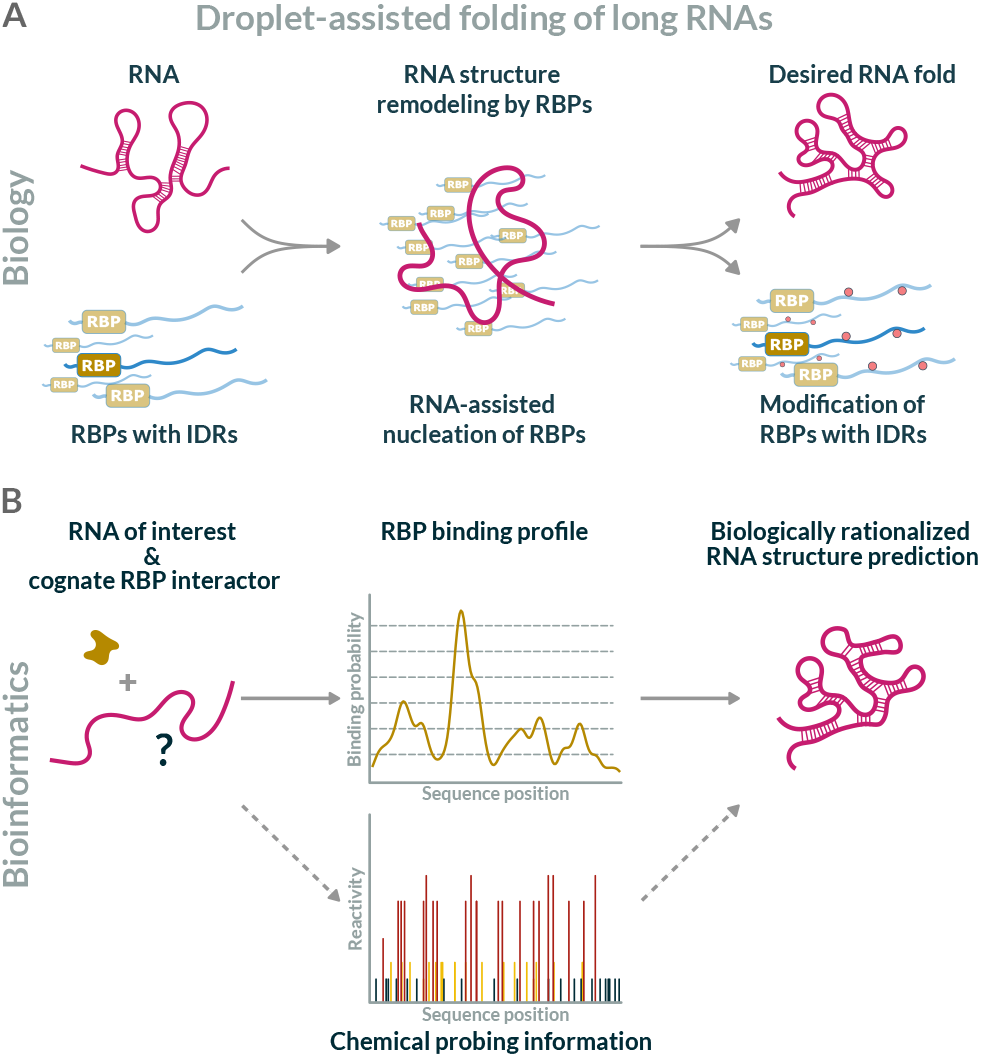
Reshaping RNA structures by condensates & mimicking the process *in silico*. **A**) RNAs and RBPS form condensates, which can remodel and dissolve the structure of RNAs. Cellular processes like post-translational modifications of RBPs can regulate and dissolve condensates. In this process, RNAs can obtain a new structure. **B**) For a given pair of RNA and cognate RBP we can calculate a binding profile. Using this, and possibly probing data, we can guide structure prediction algorithms to obtain a unique, biologically motivated RNA structure.

In this work we showed that initial cornerstones comprised of distant parts of long RNAs naturally emerge in droplet-assisted folding pathways. In contrast, traditionally considered RNA folding pathways start with cornerstones comprised of nearby nucleotides. The placement of the initial cornerstones constrain the folding trajectories and the shapes that can form; this is especially consequential for the structures of long RNAs. Droplet-assisted folding rests on the premise that as soon as phase-separating RNA-binding proteins are involved, the combination of RNA:RNA, protein:RNA, and protein:protein interactions play a critical role modulating the folding process compared to scenarios which are dominated by RNA sequence alone.

Droplets have been shown to bring distal regions of other biopolymers into close proximity: DNA enhancers can be far away from transcription sites but brought close together by droplets [82]. Double stranded DNA breaks can be kept together by PARP1-induced DNA damage repair droplets [39]. Microtubules can be bend around droplets via capillary forces bringing their distal parts together [83]. Reshaping single stranded RNA to bring distal regions into close proximity to create initial cornerstones is an extension of this logic. RNAs emerging from dissolving droplets have constrained ensembles, reducing the available conformational space and increasing the likelihood of long-range interactions [27, 45]. Thus, a droplet-assisted folding process offers a mechanistic explanation for how complex RNA structures form in vivo — something not captured by previous models.

RNA-protein interactions and the phase behavior of RBPs are highly tunable through post-translational modifications [49, 84, 85]. These modifications alter the charge distribution on RBPs, modulating their ability to bind RNA, and form droplets. As a result, droplet formation can be dynamically regulated in time and space. Interestingly, the degree of RNA binding and the degree of phase separation could be tuned independently by different enzymes [52, 86], providing additional tuning over the droplet-assisted folding trajectory.

The patterning of RBP contacts along an RNA determines the timing of detachment from different regions during droplet dissolution. Where proteins bind strongest, they detach last. Thus, when RNA regions are released for folding likely carries a sequence specificity, resulting in a narrowing of the folding landscape **Fig. 7B**).

We also envision that droplet-assisted folding can be extended to more complex scenarios, where multiple condensate droplets with distinct RBP compositions form on the same RNA. This is also supported by differences in binding density distributions of various RBPs along an RNA’s sequence (see **Fig. S6**). Such systems would better reflect the heterogeneity of protein-RNA interactions within cells and could result in alternative RNA conformations depending on the droplet’s composition and location [18, 43, 87]. Furthermore, post-transcriptional RNA modifications [88, 89] could lead to differential protein:RNA or RNA:RNA interactions affecting the folding landscape or, alternatively, the droplet-assisted folding landscape could affect the pattern of post-transcriptional modifications.

Unlike proteins, whose folding often funnels toward a single global minimum, RNA free energy landscapes are rugged, with many near-equivalent conformations [1, 8]. Droplet-assisted RNA folding provides a new lens for viewing the traversal of this complex RNA folding landscape in a physiological context.

## Methods

### Implementation of different folding algorithms: ViennaRNA, RNAstructure

To predict RNA secondary structures, we have used the Python package ViennaRNA v2.7.0 [9]. To predict structures using RNAstructure [10], we used the Julia package RNAstructure.jl, accessed through the Python module JuliaCall. Unless otherwise specified, all predictions use the standard parameters.

### Implementation of different folding scenarios

All RNA folding scenarios have been implemented using custom written Python code. For the **Full Fold** scenario the RNA sequence is passed to the folding algorithm without any restrictions. All the other scenarios are calculated iteratively with constraints being passed either directly to ViennaRNA or as a constraint file (.con extension) to RNAstructure. For the **Sequential Fold** scenario only the first N nucleotides from the 5’-end are initially accessible with the rest being forced to remain single-stranded, where N is given by the folding window. In each step, the available sequence is extended by N nucleotides, whereby the previously found base pairs are retained. This is repeated until the full sequence is accessible. The **Single Droplet** scenario follows the same principles, but initially, the first N nucleotides on both ends are accessible and increased by N more nucleotides at each side per iteration. For the **Multiple Droplet** scenario, four seeds are distributed equally over the RNA sequence. The initial droplet radius defines the nucleotides before and after each seed, which are unavailable in the initial prediction. It is typically chosen to span the whole sequence with minimal overlap. In each iteration, this distance is reduced by a droplet radius step. For the **RBPFold** scenario binding levels are derived from a binding profile (see below) using a custom Python script. Each nucleotide with a binding density below the level height is available, while nucleotides with a higher binding density are prohibited from forming base pairs. The structure is predicted iteratively with increasing levels while found base pairs are maintained. Finally, the **RBPFold + Probing** scenario is implemented in the same way, whereby the probing data is added as a soft constraint. To improve calculation speed, unavailable nucleotides are replaced by a 35-nucleotide-long poly-A sequence.

### RNA structure prediction analysis

For every RNA structure predicted by the pipeline, a series of parameters was calculated: The minimum free energy values, end-to-end distance, maximum ladder distance, and nucleotide pair distance distribution (**Figure 1** and **Figure S1**). Finally, the basepair network was also visualized as a 2D contact map, and the map was then binned into a normalized heatmap to simplify the visual interpretation (**Figure S2**). For more details, please refer to the **supplementary materials**.

Also, we have shown that the prevalence of certain structural characteristics depends mainly on the particular folding scenario and that the applied algorithm (ViennaRNA or RNAstructure) or the folding horizon only affects the resulting RNA structure to a minor degree (**Fig. S2**).

### Generation of the random, promiscuous, and scrambled RNA sequences

The random RNA sequences of a given length (2150 nt) were generated using a self-written Python script. The scrambled sequences of full-length HOTAIR RNA (2150 nt) were generated using a self-written Python script by randomly permuting the HOTAIR nucleotides.

### Extraction of the experimental RNA structures

A representative set of RNA structures was extracted from the RCSB PDB database [56]. We set the minimal length threshold to 400 nt and excluded any in vitro RNA origami structures, since we were interested in native RNA folding. To prevent bias towards certain RNA species, we selected only one example structure per specific RNA group (e.g. 16S, 18S, 23S, 25S rRNAs, Group I, II, IIC introns). We then extracted the secondary RNA structure from the PDB files using RNApdbee 2.0 [90]. For more details, please refer to the supplementary materials.

### Cloning and expression of YBX1 protein fused with mEGFP

The human YBX1 coding sequence (UniProt ID: P67809) was cloned into the pOCC127 vector using NotI and Ascl restriction sites. For insect cell expression, the insert was positioned between an N-terminal maltose-binding protein (MBP) tag with a 3C PreScission protease site and a C-terminal TEV-monoGFP-3C-His6 tag.

### Purification of YBX1 protein fused with mEGFP

Recombinant YBX1-mGFP protein was expressed in Spodoptera frugiperda (Sf9) insect cells using the baculovirus expression system. A 1 L culture of Sf9 cells at a density of 2 * 10^6^ cells/mL was transfected with recombinant baculovirus encoding the YBX1-mGFP fusion protein using a 1:50 virus-to-culture ratio. Infected cells were harvested for protein purification.

### Purification of YBX1 protein fused with mEGFP

Infected Spodoptera frugiperda (Sf9) insect cells were incubated for 64 hours at 27 °C with shaking at 100 rpm and harvested by centrifugation for 15 minutes at 1200 rpm. Cell pellets were resuspended in 60 mL lysis buffer containing 50 mM Tris-HCI (pH 8), 1 M NaCl, 5% glycerol, 10 mM imidazole, and 1 mM DTT, supplemented with 1x cOmplete ™ Protease Inhibitor Cocktail tablet (Roche, 11836170001) and 0.5 units/mL Benzonase® nuclease (Sigma-Aldrich, 9025-65-4). Cells were lysed using an LM10 Microfluidizer ™ set to 5000 psi. The lysate was clarified by centrifugation at 20,000 rpm for 60 minutes at room temperature. The supernatant was loaded onto two 5 mL Protino® Ni-NTA columns (Macherey-Nagel) in tandem for His-tagged protein purification using an ÄKTA Go ™ system (Cytiva). The column-bound protein was washed with 10 column volumes (CV) of Ni-NTA wash buffer (50 mM Tris-HCI, pH 8, 1 M NaCl, 5% glycerol, 20 mM imidazole, 1 mM DTT, 1x cOmplete T™ PIC), followed by elution using Ni-NTA elution buffer (50 mM Tris-HCl, pH 8, 1 M NaCl, 5% glycerol, 250 mM imidazole, 1 mM DTT).

Eluted peak fractions were directly applied to 4 mL amylose resin (New England Biolabs, E8021S) packed into a 20 mL gravity-flow column. The resin was washed with 8 CV of MBP/Amylose wash buffer A (50 mM Tris-HCI, pH 8, 1 M NaCl, 5% glycerol, 1 mM DTT), followed by 2 CV of MBP/Amylose wash buffer B (50 mM Tris-HCI, pH 7.8, 1 M NaCl, 5% glycerol, 1 mM DTT). MBP-tagged YBX1-mGFP fusion protein was cleaved on-column by incubating overnight at room temperature with 0.01 mg/mL 3C protease. The cleaved proteins were collected in the flow-through.

This was followed by heparin column chromatography. The cleaved YBX1-mGFP protein was diluted and equilibrated to 160 mM NaCI (pH 7.4) and loaded onto a 1 mL HiTrap T™ Heparin HP affinity column (Cytiva) connected to the ÄKTA Go ™ system. The column was washed with 8 CV of Heparin binding buffer (50 mM Tris-HCl, pH 7.4, 160 mM NaCI, 5% glycerol, 1 mM DTT), and the bound protein was eluted using a stepwise gradient of the binding buffer, increasing NaCl concentration from 160 mM to 1 M over 10 CV. Peak fractions were pooled and concentrated to 4-5 mL using a Vivaspin® 20 concentrator (30 kDa MWCO, Sartorius, VS2022). The protein was further purified by size-exclusion chromatography (SEC) using a HiLoad ™ 16/600 Superdex™ 200 pg column (Cytiva) equilibrated in storage buffer (50 mM Tris-HCI, pH 7.4, 500 mM KCI, 5% glycerol, 1 mM DTT). SEC peak fractions corresponding to the YBX1-mGFP fusion protein (65.6 kDa) were pooled, concentrated using Vivaspin® 20 (30 kDa MWCO), aliquoted (5-10 µL) into PCR tubes, snap-frozen in liquid nitrogen, and stored at −80 °C.

Protein concentration and purity were determined using a DeNovix spectrophotometer by measuring absorbance at 260, 280, and 488 nm. The 260/280 ratio was used to assess nucleic acid contamination. Extinction coefficients of 52,845 M^*−*1^ cm^*−*1^ (280 nm) and 56,000 M^*−*1^ cm^*−*1^ (488 nm) were used for calculations.

### Cloning and preparation of HOTAIR lncRNA

For IVT, HOTAIR lncRNA was purchased from Addgene (LZRS-HOTAIR, #26110) and subcloned into pCRII. PCR amplified DNA was used as a template for *in vitro* transcription with the MEGAScript T7 Transcription kit (Invitrogen), while substituting 25% of UTP with Aminoallyl-UTP-PEG5-STAR 580 (Jena Bioscience). The reaction was incubated for 3 h at 37 °C, including additional DNase treatment for 30 min. The labelled RNA was extracted with phenol/chloroform, ethanol precipitated and stored in nuclease free water at −80 °C until use.

### Spinning disc microscopy of YBX1 and HOTAIR

For colocalization, 1 nM HOTAIR-580 RNA was mixed with 3 µM YBX1-GFP in low salt buffer (50 mM Tris-HCl pH 7.4, 50 mM KCl, 0.5 mM MgCl2, 1 mM DTT, RNAseOUT 20U/ml) in 384-well PhenoPlates with glass bottom (PerkinElmer). The images were acquired after 30 minutes incubation at room temperature on an inverse Nikon Spinning Disc microscope equipped with a 100x/1.46 Oil DIC (Plan-Apochromat) objective. Images were taken in a single z-plane with sequential excitation using the 488 and 561 lasers and processed in Fiji. Scale bar = 20 µM.

### RBP binding profile calculations

For a given pair of RBP and RNA sequences, the ATtRACT database [64] containing published RBP binding motifs was searched and the published motifs were then screened against the given RNA sequence. 1 mismatch was allowed within the motif. The kdeplot function of python package seaborn was used to calculate and plot the binding site density with a bandwidth parameter = 0.2.

### Extraction of the RNA structure information for SARS-CoV-2 genomic RNA

The publicly available data by Lan *et al*. contained both the reactivity data and the full predicted structure [72]. Therefore, we used this dataset for (**i**) comparison with our predictions, (**ii**) the reactivity data as an input for our RBPFold+Probing prediction. To acquire information about long-range interactions within the SARS-CoV-2 genomic RNA, we utilized the experimental cross-linking dataset published by Ziv et al. 2020 [34]. As no full structure prediction was available, we utilized the published supplementary table, which contained all the mapped chimeric reads along with the distances between the two read parts. This distance distribution is similar to our nucleotide pair distance distribution, as it provides information about the overall distribution of distances between nucleotides within the experimental/predicted structures. Before plotting the experimental distance distribution, we filtered out all events where the read distance was negative, as these cases were predominantly enriched in the non-cross-linked controls and were therefore likely to represent an experimental artifact.

### SARS-CoV-2 Nucleocapsid protein binding profile generation

The dataset with the Nucleocapsid protein binding motif distribution published by Fan, Sun and Fan et al. 2022 was used to reconstruct the binding profile [77]. The dataset consisted of probability values of the Nucleocapsid protein binding to any given nucleotide within the SARS-CoV-2 sequence. We therefore applied a threshold of 0.001 to distinguish between the likely binding sites and the rest. The kdeplot function of python package seaborn was used to calculate and plot the binding site density with a bandwidth parameter = 0.2.

## Acknowledgements

We would like to thank Michael Schlierf, Andreas Hartmann, Manthan Raj, Matthew Kraushar, Rainer Nikolay, and Ankur Jain for several fruitful discussions on the topic. We would also like to thank Michael Schlierf and Andreas Hartmann for their feedback and critical reading of the manuscript. During parts of this work, L.P. has been supported by a Fellowship of the Peter and Traudl Engelhorn Foundation. This work has been supported by the Deutsche Forschungsgemeinschaft (DFG, German Research Foundation) under Germany’s Excellence Strategy—EXC-2068–390729961.

